# Quantification of DNA methylation using methylation-sensitive restriction enzymes and multiplex digital PCR

**DOI:** 10.1101/816744

**Authors:** R.J. Nell, D. van Steenderen, N.V. Menger, T.J. Weitering, M. Versluis, P.A. van der Velden

## Abstract

Epigenetic regulation is important in human health and disease, but the exact mechanisms remain largely enigmatic. DNA methylation represents one well-studied aspect of epigenetic regulation, but is challenging to quantify accurately. In this study, we introduce a digital approach for the absolute quantification of the amount, density and allele-specificity of DNA methylation. Combining the efficiency of methylation-sensitive restriction enzymes with the quantitative power of digital PCR, DNA methylation is measured accurately without the need to treat the DNA samples with sodium bisulphite. Moreover, as the combination of PCR amplicon and restriction enzyme is flexible, the context and density of DNA methylation can be taken into account. Additionally, by extending the experimental setup to a multiplex digital PCR, methylation markers may be analysed together with physically linked genetic markers to determine the allele-specificity of the methylation. *In-silico* simulations demonstrated the mathematical validity of the experimental setup. Next the approach was validated in a variety of healthy and malignant reference samples in the context of *RASSF1A* promotor methylation. *RASSF1A* is an established tumour suppressor gene, that is aberrantly methylated in many human cancers. A dilution series of well-characterized reference samples cross-validated the sensitivity and dynamic range of the approach. Compared to conventional PCR based methods, digital PCR provides a more accurate and more sensitive approach to quantify DNA methylation. As no sodium bisulphite conversion is needed, also analysis of minute amounts of DNA could be carried out efficiently.

## INTRODUCTION

Epigenetic gene regulation is a complex process in which multiple mechanisms are involved. DNA methylation, typically involving the reversible formation of 5-methylcytosine in a CpG context, is an important epigenetic mark in humans, that has been correlated with gene regulation in both health and disease. In cancer, aberrant DNA methylation can disturb gene expression and might mimic the effect of oncogenic mutations (1). The exact consequences of DNA methylation may be far-reaching, and heterogeneous patterns are frequently found (see **Figure 1**). A particular DNA sample may comprise multiple alleles with varying DNA methylation, and in bulk analyses these mixtures are not always correctly interpreted (2). In order to answer the questions surrounding DNA methylation and epigenetic regulation in general, accurate and quantitative DNA methylation assays are required.

**Figure 1.**
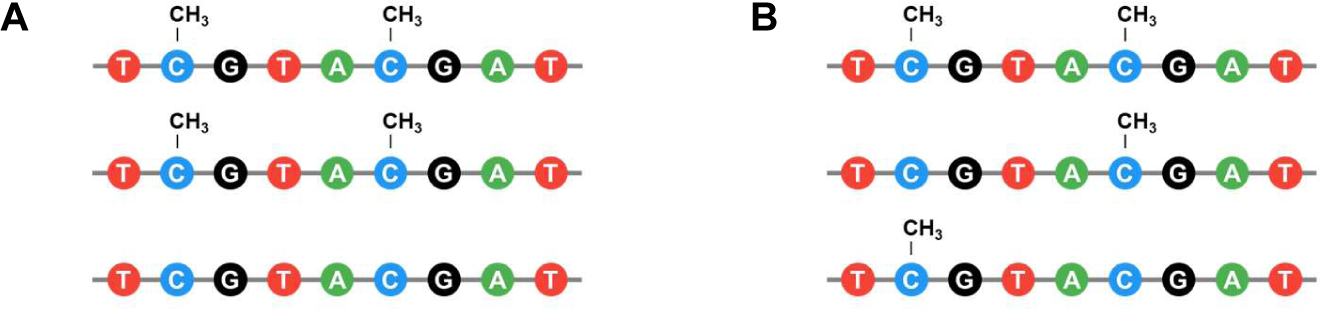
Two forms of heterogeneity in DNA methylation. **(A)** A simple mixture of methylated and unmethylated alleles. **(B)** A mixture of multiple alleles with different patterns of methylation.

Digital PCR has been proven to accurately address the genetic heterogeneity of (tumour-derived) DNA samples. Mutations, copy number alterations and, most recently, cell-type specific DNA markers have been quantified successfully to gain insight into the content and the evolutionary history of tumours (3-5). However, to measure DNA methylation in a similar fashion, new analysis methods are needed to translate available markers into quantitative assays.

In this study, we present a digital PCR-based approach to quantify a targeted density of DNA methylation through accurately measuring DNA digestion by a methylation-sensitive restriction enzyme (MSRE). Whereas most analogue methylation analysis methods depend on the chemical conversion of the input DNA with sodium bisulphite, MSRE’s can differentiate between methylated and unmethylated alleles independently of this conversion. Earlier studies already showed the potential of combining MSRE’s with qPCR or digital PCR, but were mainly focussed on the detection of low fractions methylated DNA, and on benign conditions (6-8). Instead, our approach provides a complete quantification of DNA methylation (with confidence intervals) in the whole range from 0% to 100% and can be applied in copy number unstable specimens, such as malignancies. Additionally, an extended multiplex digital PCR setup enables the combined analysis of methylation markers and physically linked genetic markers to determine the allele-specificity of the methylation.

We evaluate our approach in the context of *RASSF1A* promotor methylation. *RASSF1A* (RAS Association Domain Family 1 transcript A) is an established tumour suppressor gene that is located on chromosome 3p21, and part of several tumorigenic molecular pathways (9). Epigenetic silencing in the form of DNA promotor methylation of this gene has been demonstrated in a broad variety of human malignancies. Most notably, frequencies of up to 88-99% affected cases have been reported in lung, prostate and breast cancer (9). In healthy tissue, *RASSF1A* promotor methylation is rare and restricted to embryonic and stem-cell like cell types only making it a unique marker for foetal DNA, routinely used in prenatal diagnostic analyses (10,11).

To validate our methodological setup, an *in-silico* simulation of the digital PCR experiments is designed to verify the mathematical validity. A range of reference samples and an innovative dilution series are analysed to investigate the sensitivity and dynamic range of the approach.

## MATERIALS AND METHODS

### Sample collection and DNA isolation

5 cancer cell lines derived from primary uveal melanomas (92.1, Mel-202, Mel-270, Mel-285 and Mel-290) were available in the Department of Ophthalmology, Leiden University Medical Center, the Netherlands (12,13).

5 male PBMC (Sanquin, Amsterdam, the Netherlands) and 3 placenta samples were a kind gift from the Department of Immunohematology and Blood Transfusion, Leiden University Medical Center, the Netherlands. These anonymised samples were obtained from healthy volunteers who provided informed consent. All specimens were handled in accordance with the institutional and national ethical guidelines and the Declaration of Helsinki. DNA was isolated using the QIAmp DNA Mini Kit, according to the instructions supplied by the manufacturer (Qiagen, Hilden, Germany).

3 commercially available control DNA samples were purchased from Merck Millipore (Burlington, USA): CpGenome Universal Methylated DNA (enzymatically methylated human male genomic DNA), CpGenome Universal Unmethylated DNA vial A (human genomic DNA) and B (genomic DNA from a primary human foetal cell line).

### Experimental design

Methylation-sensitive restriction enzyme BstUI (New England Biolabs, Ipswich, USA) was selected based on its capacity to distinguish methylated from unmethylated DNA sequences. An incubation of 1 hour at 60 °C results in the digestion of unmethylated 5’-CGCG-3’ substrate sequences, whereas methylated 5’-CGCG-3’ sequences remain intact. To quantify DNA methylation, this digestion is measured in two separate duplex digital PCR experiments: one baseline experiment (referred to as *Baseline*) measuring the initial presence of target of interest, and one experiment (referred to as *MSRE+*) measuring the presence after MSRE digestion (see **Figure 2A**). To correct for input differences between the *Baseline* and *MSRE+* experiment, an independent and undigestible reference was measured simultaneously in both experiments.

**Figure 2.**
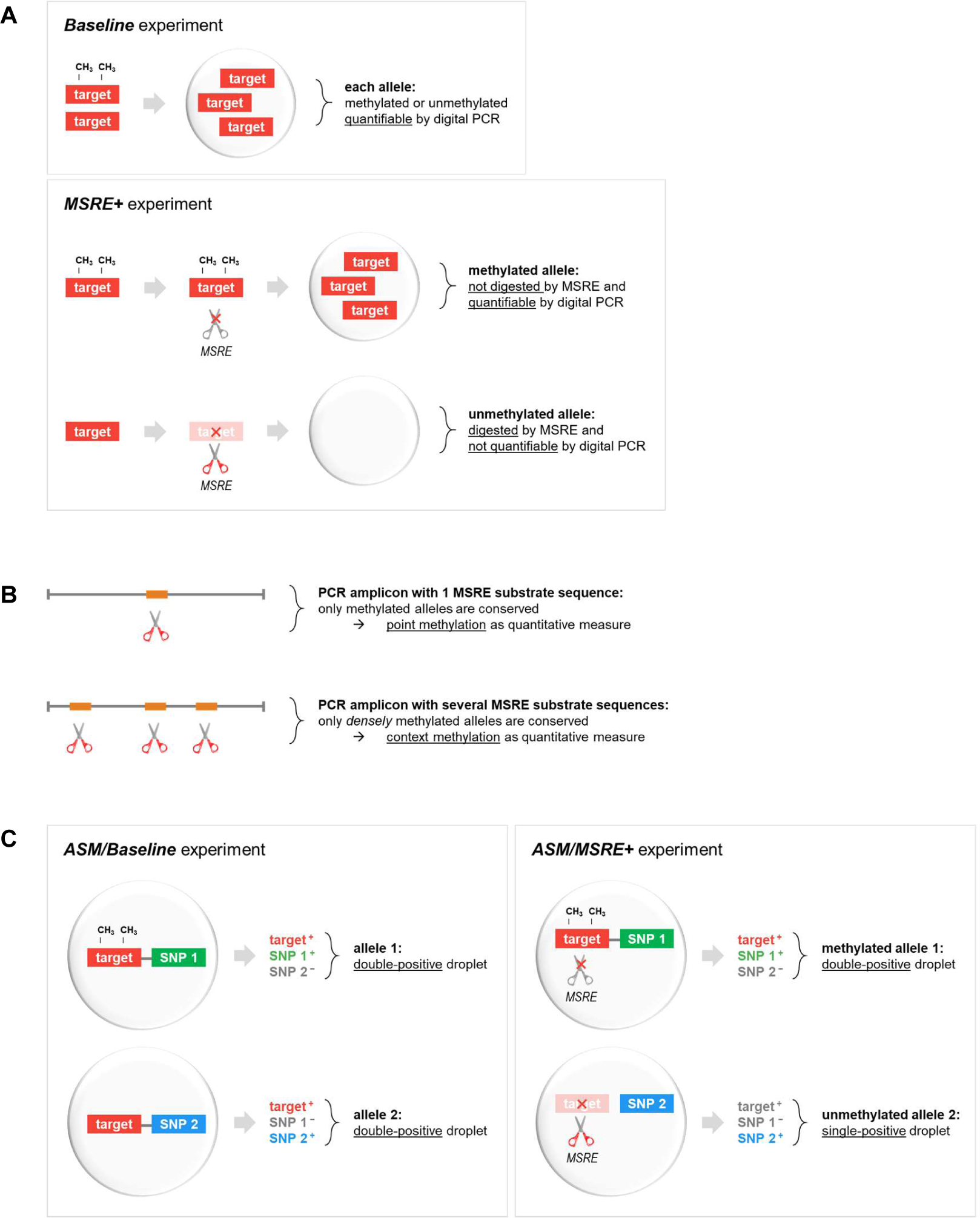
Concept of quantifying DNA methylation using MSRE and digital PCR. **(A)** A MSRE incubation of a DNA sample results in the digestion of unmethylated fragments by the MSRE, whereas methylated sequences remain intact. To calculate the fraction methylated alleles, the MSRE digestion is measured in two separate duplex digital PCR experiments: one *Baseline* experiment measuring the initial presence of target of interest, and one *MSRE+* experiment measuring the presence after MSRE digestion. **(B)** The density of DNA methylation can be integrated in the quantification by the selection of PCR amplicon and MSRE. **(C)** A methylation marker may be measured in combination with a linked genetic marker (i.e. a heterozygous SNP) to determine the allele-specificity. As intact alleles (with both the methylation target and one of the SNP variants) will end up in one digital PCR partition (i.e. droplet), a double-positive signal will be measured. After MSRE digestion within the droplet, unmethylated target will be digested, leading to single-positive signals, while methylated targets are conserved and still result in double-positive signals.

To quantify *RASSF1A* promotor methylation, a specific FAM digital PCR assay was designed (Sigma-Aldrich, Gillingham, Dorset, UK) targeting the promotor region of *RASSF1A* and five 5’-CGCG-3’ BstUI recognition sequences. Only alleles with consecutive methylation on these sequences are conserved by MSRE treatment and measurable in the *MSRE+* experiment. In contrast, (partly) unmethylated alleles are digested by MSRE treatment. As independent stable reference, a HEX digital PCR assay for *TERT* was used (Bio-Rad, Hercules, USA), in which no BstUI recognition sequences were present.

The presence of *RASSF1A* is compared against reference *TERT* in each experiment by calculating the ratio *RASSF1A/TERT*:

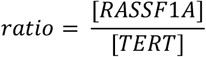

The methylation fraction (defined as the fraction methylated *RASSF1A* alleles of all *RASSF1A* alleles) is calculated by dividing the ratios *RASSF1A/TERT* of the *MSRE+* and *Baseline* experiment:

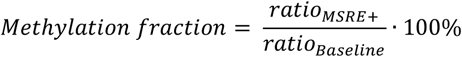

Note that in copy number stable samples the concentration of all *RASSF1A* alleles can be estimated by the concentration of *TERT* alleles. The ratio *RASSF1A/TERT* of the *MSRE+* experiment then already gives the methylation fraction. For consistency across all experiments, this approach was not applied in any of our samples.

In copy number unstable samples, *TERT* might be involved in genetic alterations. As *TERT* is used to normalise input differences between the *MSRE*+ and *Baseline* experiment, its only requirement is being measurable without being affected by the MSRE. Therefore, a TERT copy number alteration would generally not influence the methylation fraction of RASSF1A.

Confidence intervals on obtained concentration ratios can be calculated using the geometric interpretation of Fieller’s theorem (14). This mathematical approach could easily be translated to our experimental design, as outlined in **Supplementary data 1**. Following these formulas, an indication of the subsampling error in our methylation quantifications can be calculated at a given level of significance.

For the *in-silico* simulations, *R* (version 3.6.0), *RStudio* (version 1.1.463) and R packages *rmarkdown* (version 1.11) and *digitalPCRsimulations* (version 1.1.0) were used.

MIQE context sequences for *RASSF1A* and *TERT* digital PCR assays are presented in **Supplementary table 1**.

### MSRE incubation and digital PCR

Generally, 50 ng of DNA was incubated for 1 hour at 60 °C with 1 U BstUI and 0.5 uL 10x CutSmart buffer (New England Biolabs), in a total volume of 5.0 uL. For *Baseline* experiments, 50 ng of DNA was incubated for 1 hour at 60 °C with 0.5 uL 10x CutSmart buffer, in a total volume of 5.0 uL. All incubations took place in a T100 Thermal Cycler (Bio-Rad Laboratories).

Digital PCR experiments were performed using the QX200™ Droplet Digital™ PCR System (Bio-Rad Laboratories) following the general experimental guidelines as described extensively by Zoutman et al. (15). In short, 20 ng of incubated DNA was analysed in a 22 uL experiment, using 11 uL ddPCR™ Supermix for Probes (No dUTP, Bio-Rad Laboratories) and primers and probes in a final concentration of 900 and 250 nM respectively.

PCR mixtures were partitioned into 20.000 droplets using the AutoDG™ System (Bio-Rad Laboratories). Subsequent PCR was performed in a T100 Thermal Cycler using the following protocol: 10 min at 95 °C; 30 s at 94 °C and 1 min at 55 °C for 40 cycles; 10 min at 98 °C; cooling at 12 °C for up to 48 h, until droplet reading. Ramp rate was set to 2 °C/s for all steps.

Droplet reading was performed in a QX200™ Droplet Reader (Bio-Rad Laboratories).

### Determination of allele-specificity

To quantify the allele-specificity of the *RASSF1A* promotor methylation (allele specific methylation, ASM), the *TERT* reference assay was replaced by a FAM/HEX digital PCR assay for the two variants of SNP *rs1989839* (see **Figure 2C**), in which no BstUI substrate sequences were present. The sum of concentrations of both variants was taken as stable reference in calculating the ratio.

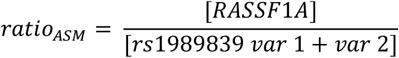

Whereas in the first experimental setup the reference was not located in the *RASSF1A* target region, this is the case for SNP *rs1989839 (∼1kb upstream)*. Assuming local copy number alterations are not present within this small region, the ratio should equal 1 in the *ASM/Baseline* experiment, and equals the methylation fraction in the *ASM/MSRE+* experiment.

MIQE context sequence for the SNP *rs1989839* digital PCR assays (Bio-Rad) is presented in **Supplementary table 1**.

The linkage between the *RASSF1A* target and the SNP variants was calculated as outlined in **Supplementary data 2**.

As the allele-specificity of the methylation will be determined based on the linkage of *RASSF1A* target and SNP variants on intact alleles, the MSRE should be prevented from fragmenting alleles between both loci. Therefore, an adapted protocol was followed in which the MSRE incubation took place after droplet generation. All experiments performed according to this protocol are marked with ‘ASM’ (allele-specific methylation).

PCR mixtures of 22 uL were prepared on ice and contained 20 ng of untreated DNA, 11 uL ddPCR™ Supermix for Probes (No dUTP) and primers and probes in a final concentration of 900 and 250 nM. For *ASM/MSRE+* experiments, 1 U BstUI and 0.5 uL 10x CutSmart buffer were added to the mixtures. For *ASM/Baseline* experiments, only 0.5 uL 10x CutSmart buffer was added to the mixtures.

To inhibit MSRE activity before partitioning into the droplets, droplet generation was performed manually using a cooled QX200™ Droplet Generator (Bio-Rad Laboratories). A MSRE incubation step of 1 hour at 60 °C was added as first step prior to the PCR protocol described earlier and carried out in one run in a T100 Thermal Cycler. Droplet reading was performed in a QX200™ Droplet Reader.

### Data analysis

Raw digital PCR results were acquired using *QuantaSoft* (version 1.7.4, Bio-Rad Laboratories) and imported in online digital PCR management and analysis application *Roodcom WebAnalysis* (version 1.8.0, https://www.roodcom.nl/webanalysis).

### Methylation-sensitive melting curve analysis and bisulphite sequencing

To validate obtained methylation fractions by methylation-sensitive melting curve analysis (MS-MCA) and bisulphite sequencing (BS), a sodium bisulphite conversion of *MSRE+* and *Baseline* samples was carried out using the EZ DNA Methylation Kit (Zymo Research Corporation, Irvine, USA). Generally, 100 ng input DNA was converted according to the manufacturer’s guidelines and eluted in 10 uL M-Elution Buffer. To remove possibly interfering salts, a DNA purification prior to the conversion was performed using DNA Clean & Concentrator (Zymo Research Corporation), following the manufacturer’s instructions.

1 uL bisulphite converted DNA was analysed in 7 uL experiment, using 3.5 uL iQ SYBR Green Supermix (Bio-Rad Laboratories) and primers in a final concentration of 700 nM. Consequent PCR was performed in a CFX384 Real-Time PCR system (Bio-Rad Laboratories), using the following protocol: 3 min at 95 °C; 10 s at 96 °C, 10 s at 65 °C and 30 s at 72 °C for 8 cycles; 10 s at 96 °C, 10 s at 62 °C and 30 s at 72 °C for 32 cycles; 10 s at 95 °C; melting curve from 65 °C to 95 °C, per 0.3 °C.

Raw PCR products were purified and analysed for ‘Quick Shot Short GC’ Sanger Sequencing at BaseClear, Leiden, the Netherlands. AB1-files were analysed using *Roodcom SangerSeq Analysis* (version 1.0, https://www.roodcom.nl/sangerseq).

MIQE context sequence for the used primers is presented in **Supplementary table 1**.

## RESULTS

### Dual-duplex digital PCR experimental setup to quantify DNA methylation

To determine the methylation fraction in a sample of interest, two separate duplex digital PCR experiments were carried out quantifying target and reference sequences before and after incubation with a MSRE. In the *Baseline* experiment all alleles (both methylated and unmethylated) were measured, while after digestion by the MSRE (i.e. the *MSRE+* experiment), only the methylated alleles were quantitated (see **Figure 2A**). The results of both experiments were compared to get a properly normalised methylation fraction.

We chose to evaluate our approach by quantifying dense *RASSF1A* promotor methylation. By selecting a PCR target amplicon containing multiple recognition sequences for the preferred MSRE, only methylation of all these so-called restriction sites (i.e. a densely methylated allele) can prevent MSRE digestion. Conceptually, this illustrates how the context of DNA methylation can be integrated in the quantification (see **Figure 2B**). As the methylation of surrounding CpG’s may be biologically relevant, a valuable aspect is included into the analysis.

### In-silico evaluation of experimental setup and mathematical rationale

The digital nature of our experimental setup enables an advanced *in-silico* modelling of it, which is described in **Supplementary data 1**. In short; for each of 50 different conditions (5 input concentrations [5, 10, 20, 50 and 100 ng genomic DNA] with 10 input methylation fractions [10%-100% in steps of 10%]), 10,000 digital PCR simulations were performed. This allowed an extensive evaluation of our experimental approach, illustrating the theoretical accuracy and precision of the measurements, and the meaning of the calculated confidence intervals.

Generally, the methylation fraction is correctly estimated across all simulated conditions (see **Figure 3A**). Though, the absolute uncertainty of the measurements becomes larger at lower input concentrations and higher methylation fractions (see **Figure 3B**). Overall, in 95.1% of the total of 1,000,000 simulations the true methylation fraction did lie in the calculated confidence interval. As this coverage is very close to our intended 95%, it shows the validity of the mathematical approach. The coverage per condition was very comparable between all input states (see **Figure 3C**). In contrast, the absolute width of the calculated confidence intervals was highly variable among amount of input DNA and input methylation fraction (see **Figure 3D**). However, as the digital error can be correctly estimated, broader confidence intervals were observed when a larger uncertainty was found, which was at lower input amounts and at higher methylation fractions.

**Figure 3.**
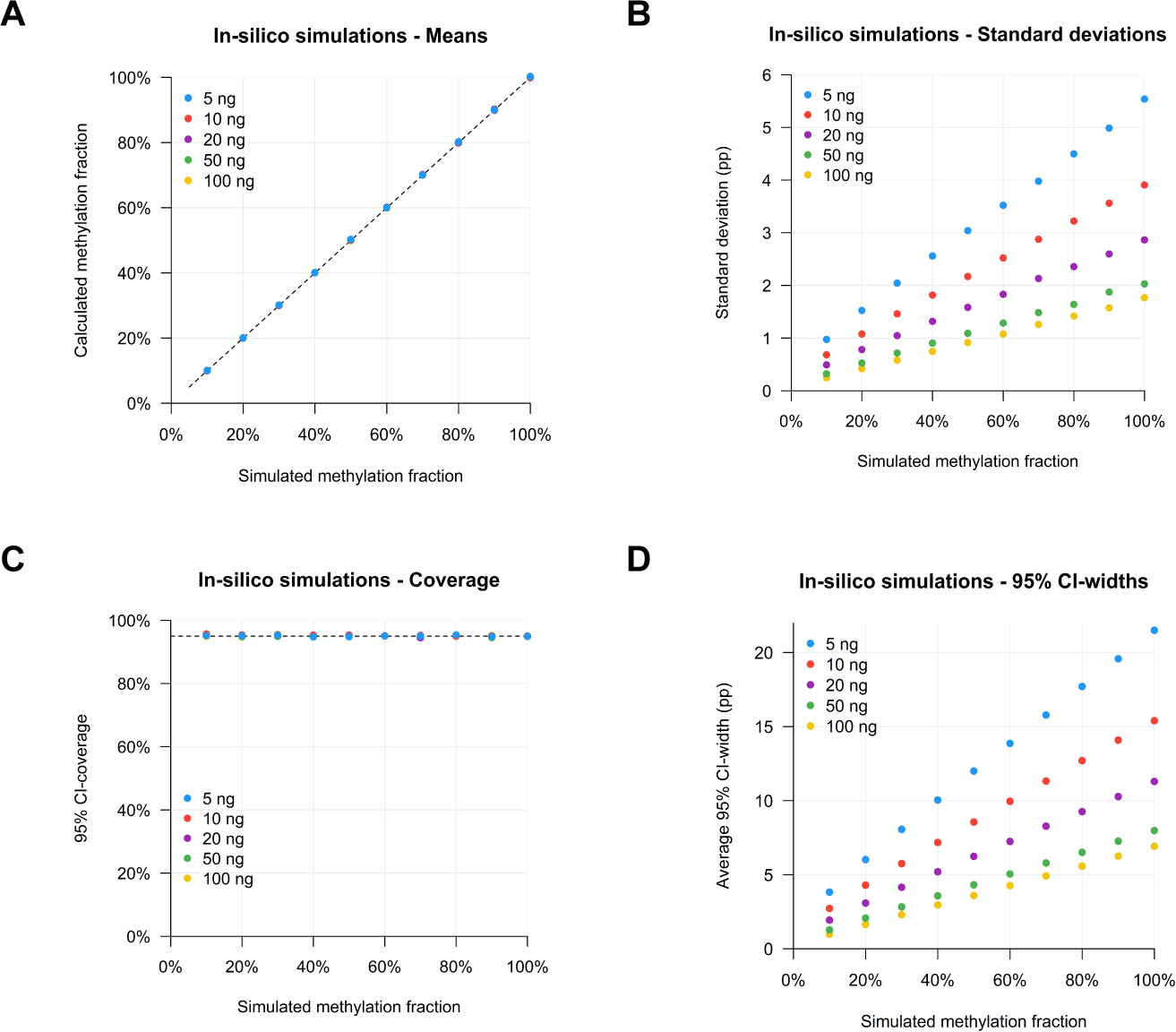
Illustration of digital approach by *in-silico* simulations for 20 methylation fractions and 5 input amount conditions. **(A)** Mean of calculated methylation fractions per input condition. **(B)** Standard deviation of calculated methylation fractions per input condition. **(C)** 95% confidence interval coverage (i.e. in which % of the simulations the true methylation fraction did lie in the calculated confidence interval) per input condition. **(D)** Average absolute widths of 95% confidence intervals per input condition.

### RASSF1A promotor methylation in reference samples

A positive biological control for *RASSF1A* promotor methylation can be found in malignancies and in embryonic tissue (10,16). Therefore, 5 uveal melanoma cell lines and 3 placental DNA samples were analysed as reference samples. An illustration of the experimental and analytic workflow to obtain the results is given in **Figure 4A**.

**Figure 4.**
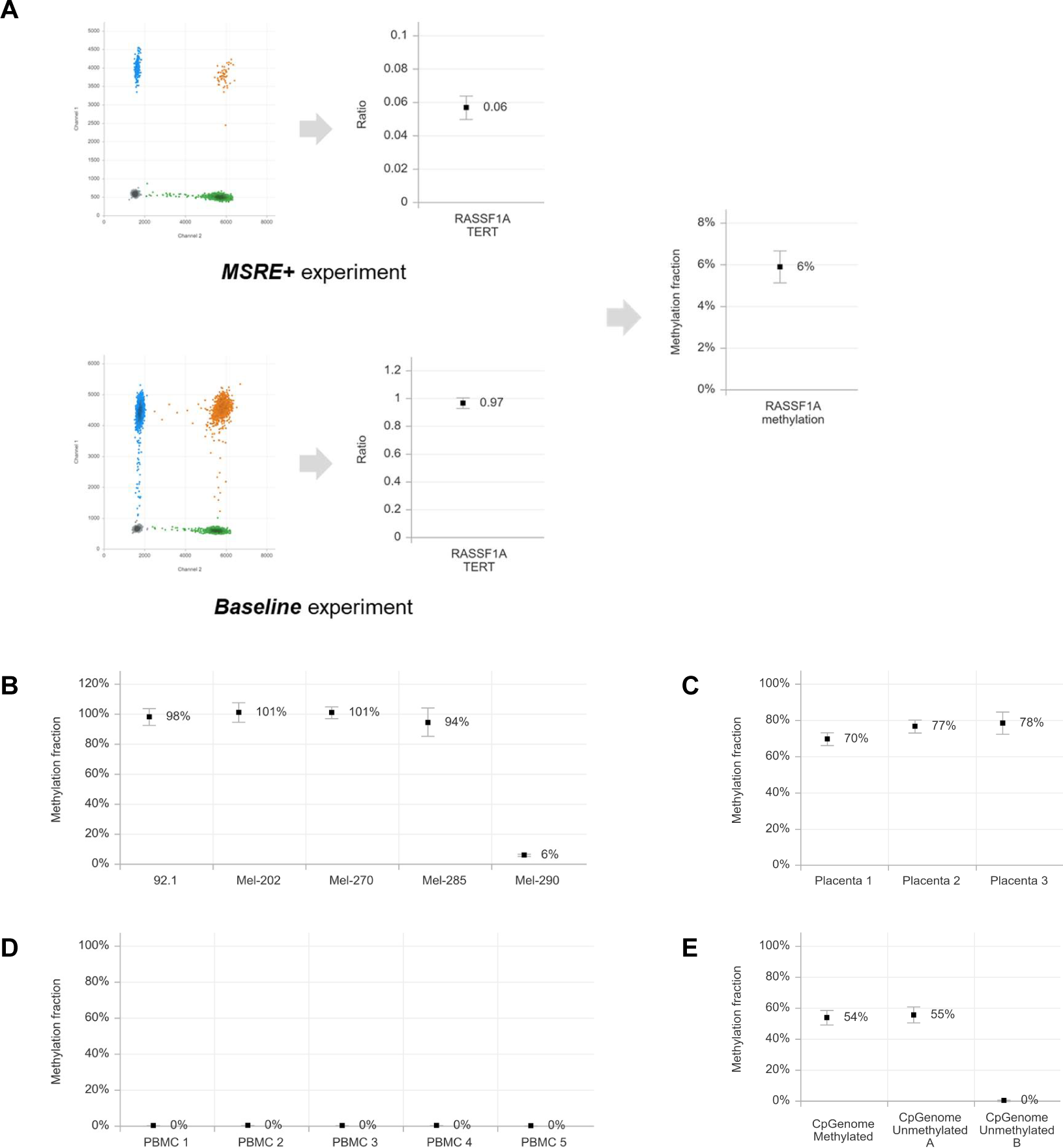
*RASSF1A* promotor methylation fractions in control samples. (**A**) Illustration of the experimental and analytic workflow to obtain the methylation fraction, example for cell line Mel-290. Summary of RASSF1A promotor methylation fractions in (**B**) 5 uveal melanoma cell lines, (**C**) 3 placenta samples, (**D**) 5 healthy male PBMC samples and (**E**) 3 CpGenome control samples.

In 4 out of the 5 uveal melanoma cell lines, in which previously extensive *RASSF1A* promotor methylation was observed, only completely methylated alleles were detected (**Figure 4B**) (17). The 5th cell line (Mel-290), the only one that was previously typed as unmethylated, now presented with 6% methylation.

In placental tissue both methylated and unmethylated cells are present due to a mix of foetal and maternal cells (10). 3 placenta DNA samples were analysed and as expected substantial *RASSF1A* promotor methylation fractions of 70%, 77% and 78% were found (**Figure 4C**).

Although in healthy female blood small amounts of methylated *RASSF1A* may be found (derived from current or prior pregnancy), healthy male blood only contains unmethylated *RASSF1A* alleles and can therefore be used as a negative control (11). Indeed, in 5 PBMC samples from healthy males, 0% *RASSF1A* methylation was detected (**Figure 4D**).

Unmethylated and methylated genomic DNA is commercially available and an attractive source of control DNA for gene methylation studies. However, in 2 of the 3 available samples partial *RASSF1A* promotor methylation was detected in our analyses. CpGenome Universal Methylated DNA demonstrated an 54% methylation fraction, whereas CpGenome Universal Unmethylated A and B presented with fractions of 55% and 0% *RASSF1A* methylation respectively (**Figure 4E**).

### Validation by MS-MCA and bisulphite sequencing

To validate our findings, a selection of samples was treated with sodium bisulphite and analysed using methylation-sensitive melting curve analysis (MS-MCA) and bisulphite sequencing (BS) (see **Figure 5 and S1**). In order to validate that only methylated alleles resisted MSRE digestion, samples that were pre-incubated with the restriction enzyme (*MSRE+*) were also included in these analyses.

**Figure 5.**
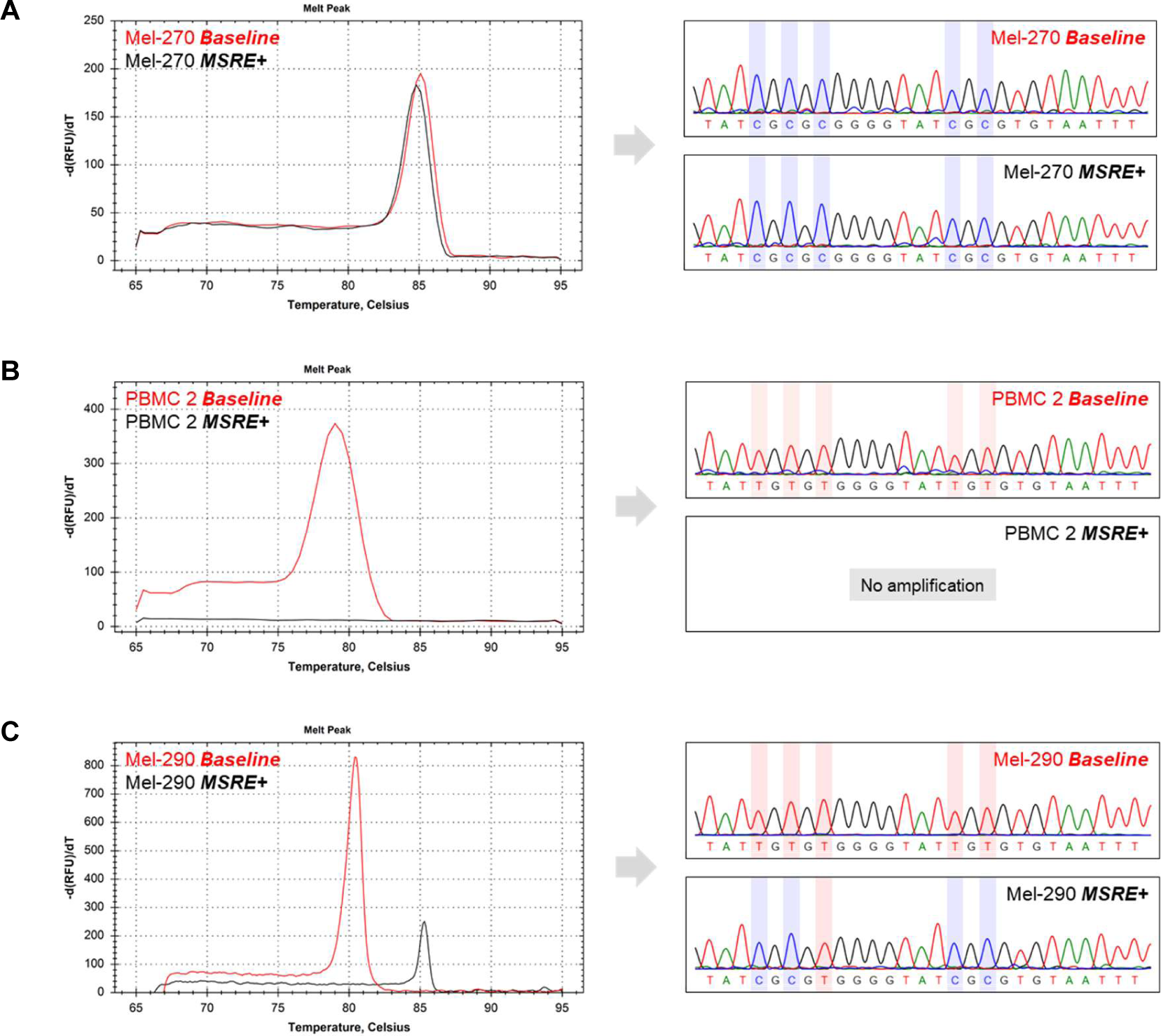
Validation of *RASSF1A* methylation in reference samples with MS-MCA and BS. **(A)** Fully methylated cell line Mel-270 presents with melting curves peaking around 85 °C, indicative of methylation. In the sequencing methylated CpG’s are found. **(B)** Fully unmethylated blood sample PBMC 2 only shows unmethylated signals (melting peak around 80 °C and unmethylated sequencing) in the *Baseline* experiment, leading to complete MSRE digestion and loss of all signal in the *MSRE+* experiment. **(C)** Only unmethylated *RASSF1A* was detected in baseline MS-MCA and BS analyses with Mel-290, while after MSRE digestion an unquantifiable methylated fraction emerged.

The fully methylated cell lines (e.g. Mel-270) were resistant to *MSRE* digestion and therefore showed similar results between the *Baseline* and *MSRE+* experiments, indicative of methylation (see **Figure 5A**).

The unmethylated blood samples (e.g. PBMC 2) did not show signs of methylation and as a consequence only generated signals in the *Baseline* experiments. Typically, the lack of methylated alleles led to a full sample digestion in the *MSRE+* experiments, resulting in no amplification in the MS-MCA and BS (see **Figure 5B**).

Regarding cell line Mel-290 (6% methylation), no signs of methylation were observed in both the MS-MCA and BS *Baseline* results. Only after the MSRE-based enrichment for methylated alleles, the presence of a small number of methylated alleles was revealed by the combination of MS-MCA or BS and MSRE enrichment for methylation (see **Figure 5C**).

The placenta samples and 2 of the 3 commercially available methylation control samples (CpGenome Universal Methylated and Unmethylated A) were all characterized by partial *RASSF1A* methylation, in which both methylated and unmethylated alleles were present. These findings were confirmed by the MS-MCA and BS results, in which in the *Baseline* experiments heterogenous peaks were found, and methylated peaks only in the *MSRE+* experiments (see **Figure S1A-D**).

### High accuracy and dynamic range

To evaluate the accuracy and dynamic range of the approach, a calibration curve was prepared. In different ratios fully methylated target DNA (derived from cell line Mel-270) was mixed with fully unmethylated DNA (derived from PBMC 2), maintaining a constant total DNA concentration. Subsequently, all mixtures were analysed for *RASSF1A* methylation. In addition, the genotype status of single nucleotide polymorphism (SNP) *rs1989839* was quantified in all mixtures. This SNP-based approach allowed us to ascertain the actual diluted methylation fraction. As samples Mel-270 and PBMC 2 can be distinguished with this SNP, mixtures of both samples can be reconstituted based on the exact presence of both alleles, see **Figure 6A**.

**Figure 6.**
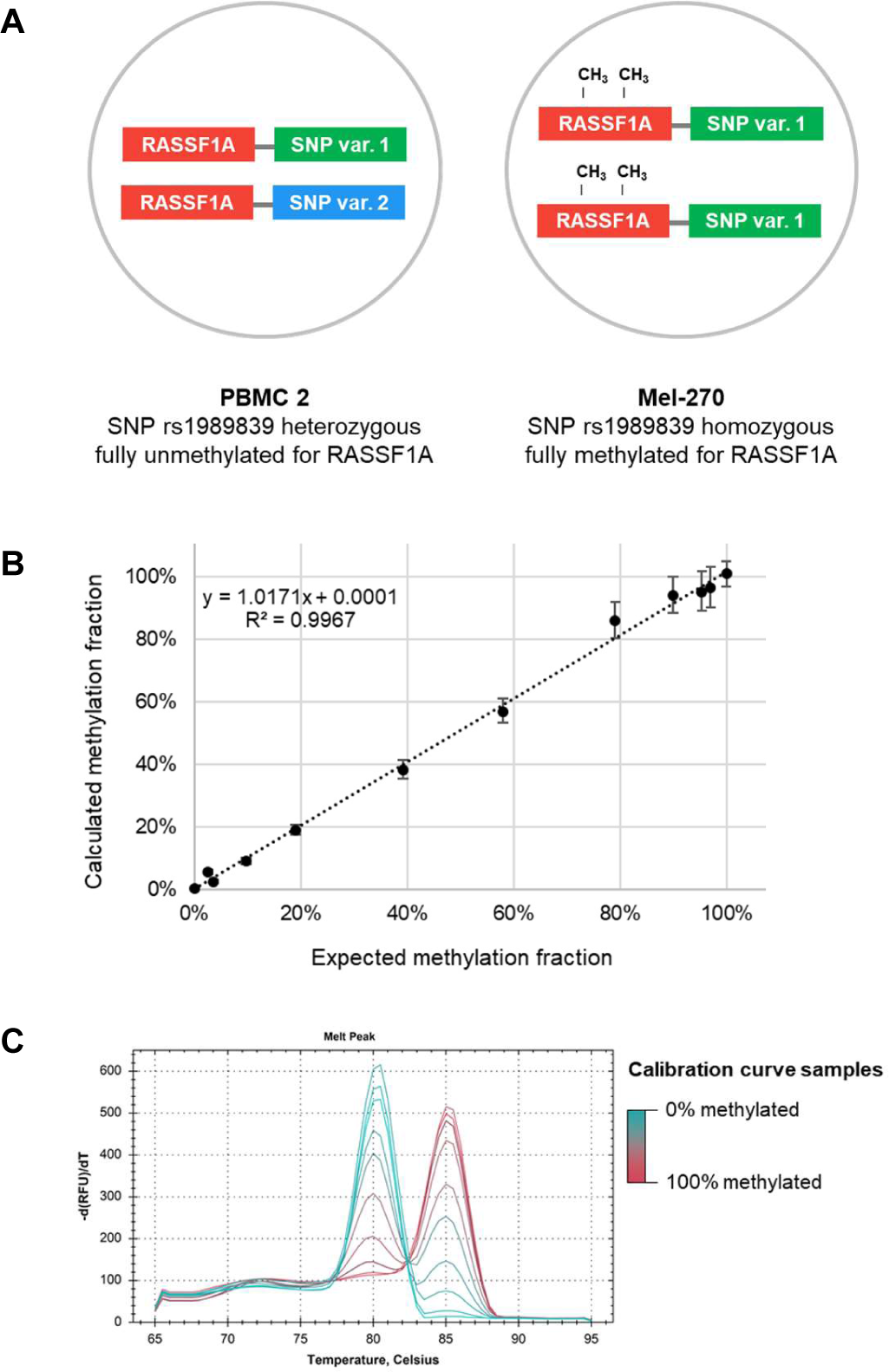
A calibration curve for *RASSF1A* methylation with diluted reference samples that includes a genetic calibrator by way of a SNP. **(A)** Equal concentrations of PBMC 2 (fully unmethylated for *RASSF1A* and heterozygous for SNP rs1989839) and Mel-270 (fully methylated for *RASSF1A* and homozygous for SNP rs1989839) are mixed in different ratios, which are measured for *RASSF1A* methylation fraction (y-axis) and SNP variant balance (x-axis). This balance, calculated as % Mel-270 = 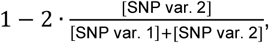, validates the mixed ratio of both samples. **(B)** A calibration curve comparing the expected and measured methylation fraction showed a high linear correlation (R^2^ = 0.9967), indicating the quantitative nature of the approach. **(C)** MS-MCA of all prepared ratios showed gradually in- and decreasing peak heights at the methylated and unmethylated amplicon melting temperatures, but could not easily be translated into a quantitative measure.

The results of both measurements are integrated in **Figure 6B**. A linear relationship between input ratio and calculated methylation fraction was observed (Pearson R^2^=0.9967, slope = 1.0171), indicative of high accuracy of the measurements.

The calibration curve samples were also treated with sodium bisulphite and analysed using MS-MCA. As expected, gradually in- and decreasing peak heights at the methylated and unmethylated amplicon melting temperatures were observed across the samples (see **Figure 6C**). This peak distribution could however not easily be translated into a quantitative value, especially as small peaks were not consistently identifiable at the lowest and highest methylation fractions.

### Allele-specific methylation

In addition to the described approach to quantify a targeted density of DNA methylation, a methylation marker may be measured in combination with a linked genetic marker to determine the allele-specificity. In our experiments, the DNA methylation target was analysed together with a heterozygous SNP nearby this target, which serves as and replaces the stable reference. These analyses were carried out in a multiplex digital PCR experiment. As intact alleles (with both the methylation target and the SNP variant) will end up in one digital PCR partition (i.e. droplet), a double-positive signal will be measured. After MSRE digestion within the droplet, unmethylated target will be digested, leading to single-positive signal for the SNP variant only, while methylated targets are conserved and still result in double-positive signals (see **Figure 2C**).

Given the 2D plot of a multiplex digital PCR experiment, the distribution of droplets into the clusters indicates the presence of non-randomly linked targets. As a quantitative measure, we propose to calculate the concentrations of unlinked SNP variant (see **Supplementary data 2**). The concentrations of linked SNP variant 1 and 2 (i.e. the fraction alleles of the SNP target that is linked to another target) can be subsequently estimated by subtracting the unlinked concentration SNP variant from the total concentration SNP variant. Similarly, the linkage for the DNA methylation target can be determined. As the sum of total concentrations of the SNP variants remains stable and thus represent the total number of analysed alleles, all values can be normalised adequately. The complete mathematical rationale is outlined and validated in **Supplementary data 2**.

Applying this approach, an *ASM/Baseline* experiment (i.e. without MSRE) will measure the initial distribution of SNP variants and concordant linkage in the sample of interest. Under the hypothetical conditions of a perfect heterozygous sample with intact alleles only, a complete dichotomy may be expected **(Figure 7A)**. Here, no single-positive droplets are present, as droplets could contain either no target allele (negative for *RASSF1A* and the SNP variants), or intact allele(s) (double-positive for *RASSF1A* and a SNP variant).

**Figure 7.**
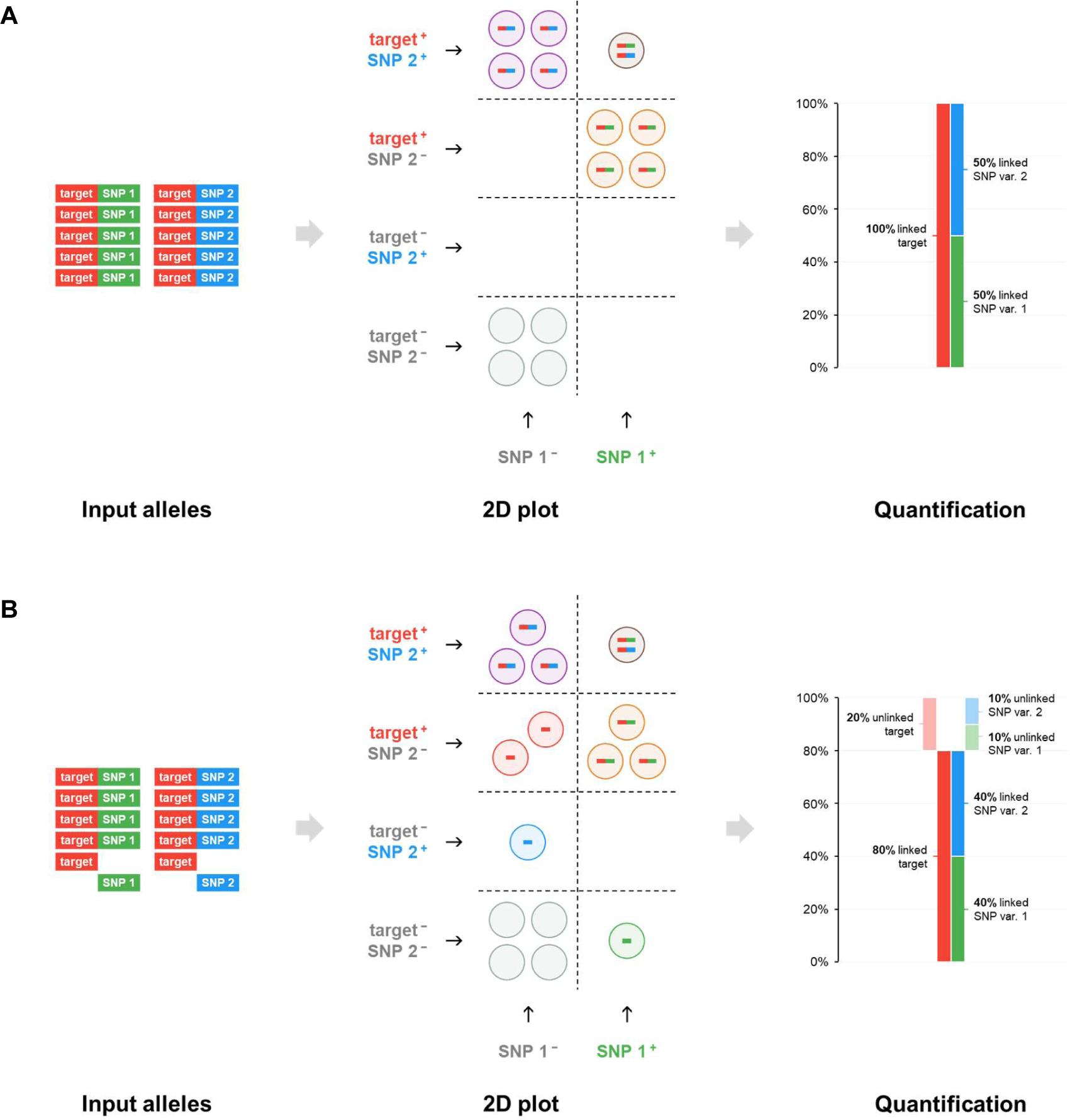
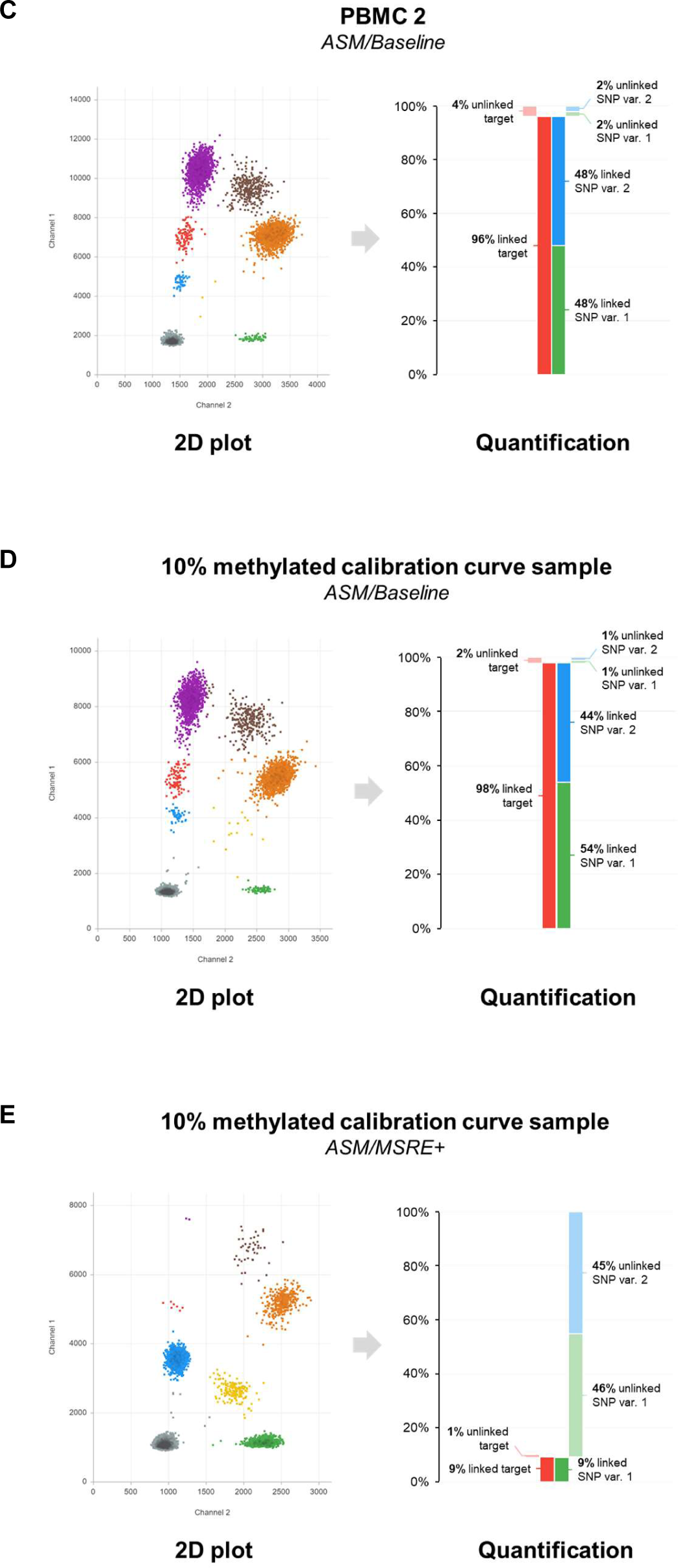
Illustrations of input conditions, 2D plots and results for *ASM* experiments. **(A)** Basic concept of ASM analysis using multiplex digital PCR in a hypothetical example with complete linkage between the heterozygous SNP and the methylation target. **(B)** A hypothetical example with 80% balanced linkage between the heterozygous SNP and the methylation target. **(C)** *ASM/Baseline* experiment without MSRE of a PBMC control sample showing 96% balanced linkage between the heterozygous SNP and the methylation target *RASSF1A*. **(D)** *ASM/Baseline* experiment without MSRE of the 10% methylated calibration curve sample showing an imbalanced presence of both SNP variants, resulting from the mixture of 10% Mel-270 (SNP var. 1 homozygous) and 90% PBMC 2 (heterozygous for SNP var. 1 and 2). Overall, a high linkage of 98% exists between all SNP variants and methylation target *RASSF1A*. **(E)** *ASM/MSRE+* experiment analysing the allele-specificity of *RASSF1A* methylation in the 10% methylated calibration curve sample. As in this sample methylation is only derived from Mel-270 (SNP var. 1 homozygous), a high allele-specific linkage is observed, with 9% of the alleles being linked to SNP var. 1 and 0% to SNP var. 2.

In a biological sample however, a certain fraction of the alleles may have been fragmented before partitioning into the droplets. We propose that e.g. the distance in nucleotides between the methylation target and the SNP, the DNA integrity, and the experimental conditions could influence this initial – and probably irreversible – loss of intact alleles. As a consequence, single-positive droplets will appear in the 2D plot and the overall linkage will be calculated being lower than 100% (see **Figure 7B**).

This proposed *ASM/Baseline* experiment was performed on a healthy, heterozygous PBMC control DNA sample (see **Figure 7C)**. Indeed, the concentrations of both SNP variants sum up to the total concentration *RASSF1A*, and heterozygosity is reflected by the 50% and 50% ratios of the SNP variants. This sample presents with limited DNA degradation as both SNP alleles are equally highly linked (48%) to the *RASSF1A* locus.

When a MSRE incubation is applied to a partitioned sample (i.e. the MSRE incubation taking place within the droplets), this so-called *ASM/MSRE+* experiment specifies the measurements to the methylated alleles (see **Figure 2C**). Whereas the original distribution of SNP variants should remain the same (the SNP amplicons cannot be digested by the MSRE), the total amount of target alleles may be lower (unmethylated target will be digested). We evaluated this extended setup in 5 of our prepared calibration curve samples, of which one is presented here. As these mixtures are generated from fully methylated Mel-270 (SNP var. 1 homozygous) and fully unmethylated PBMC 2 (SNP var. 1/2 heterozygous), *RASSF1A* methylation is only present on alleles carrying SNP var. 1.

In the 10% mixture, the *ASM/Baseline* experiment showed a clear and explainable SNP imbalance (see **Figure 7D**). As in this sample 10% homozygous SNP variant 1 DNA from Mel-270 was mixed with 90% heterozygous DNA from PBMC 2 (i.e. 45% SNP variant 1 and 45% SNP variant 2), a total presence of 55% SNP variant 1 was correctly measured. Again, both SNP variants sum up to the total amount of *RASSF1A* target and a very limited fraction of the alleles was fragmented (only 1% per SNP variant was unlinked). In the *ASM/MSRE+* experiment of this sample indeed 10% of the *RASSF1A* target alleles was methylated and persisted after the MSRE digestion (see **Figure 7E**). Additionally, a high allele-specific linkage was observed, as 9% of the alleles was linked to SNP var. 1 and 0% to SNP var. 2.

## DISCUSSION

DNA methylation, one of the main markers of epigenetic regulation, is a fundamental process involved in human embryology, physiology and pathology. Especially in cancer, aberrant methylation plays a role in the initiation and progression of disease (1).

Along with genetic alterations, also epigenetic changes diversify tumour cell populations (‘intratumour heterogeneity’). Epigenetic subpopulations may arise during tumour evolution, or due to epigenetically defined differentiation programs in normal cells (18). Recent technological advances have enabled extensive, quantitative studies into the genetically determined tumour heterogeneity, which provided new insights into the content, but also evolutionary history of tumours (4,19). However, comparable quantitative analyses for epigenetic alterations are scarce, and quantifying epigenetic heterogeneity is still challenging (2).

Common methylation analyses do not recognize the presence of epigenetic heterogeneity and produce a dichotomous outcome, comparable to the traditional genetic analyses. In this way, methylation state is interpreted as ‘yes’ or ‘no’ while the real dynamic range may be much wider (**Figure 1**). Hence, heterogeneous methylation patterns are often not correctly recognised (2).

Considering the broad variety of new applications with digital PCR-based genetic analyses, we aimed to develop a solid, comparable method to quantify epigenetic heterogeneity using digital PCR [1-3]. In this study we showed that digital PCR in combination with a MSRE allows to quantify a targeted density of DNA methylation accurately.

Widely-used methods (methylation-specific PCR, MS-MCA, BS, pyrosequencing, bead microarray) require a sodium bisulphite conversion of the DNA to make DNA methylation analysable (20). Ideally, this conversion deaminates unmethylated cytosines to uracils, but does not change methylated cytosines. However, the experimental conditions for this conversion are critical. To avoid incomplete conversion and false-positive results, an experiment is carried out that may result in extensive degradation (of up to 90%) of the input DNA (21,22). Moreover, both consistent and inconsistent conversion introduce sequence differences and hence amplification biases are not uncommon, possibly leading to incorrect quantitative interpretations (23).

In this study we prepared mixtures of fully methylated and fully unmethylated alleles, and analysed them by using our combination of MSRE and digital PCR (see **Figure 6B**). Generally, these in vitro generated mixtures validated the quantitative nature of MSRE in combination with digital PCR. Validation with MS-MCA showed patterns that were fully in line with the dilutions but nevertheless allowed only qualitative interpretation. Sensitivity also turned out to be critical in MS-MCA as the minor presence of methylated alleles in our 2.5% and 5% mixtures was undetectable, and could therefore not be quantified (see **Figure 6C**). Our digital approach, in contrast, distinguished these minor fractions of methylated alleles and showed significant results with dilutions of 2% and 5%. Similarly, uveal melanoma cell line Mel-290 presented with a small fraction of 6% methylated alleles using digital PCR, which was again undetectable with MS-MCA and BS (see **Figure 5C**) (17).

Whereas sodium bisulphite conversion incorporates the original methylation state in the converted sequence, our approach relies on the specificity and sensitivity of the chosen MSRE to distinguish methylated from unmethylated sites. Consequently, when MSRE treatment leads to non-specific or incomplete DNA digestion, under- or overestimated methylated fractions will be obtained (8). However, a calibration curve made with validated control samples indicated a properly working and highly efficient MSRE, as the calculated methylation fractions accurately matched with the full range of prepared ratios (R^2^ = 0.9699, **Figure 6B**). Importantly, as this result was obtained without a cumbersome sodium bisulphite conversion, an accurate quantification can be obtained in a faster, easier and less sample-requiring experiment.

In a methodologically comparable approach combining MSRE digestion with real-time PCR has been applied previously. Importantly, commercially available methylated and unmethylated control DNA samples were added as reference samples to the analysis in order to correct for non-specific and incomplete DNA digestion (8). However, here we showed that absolute quantification of methylated RASSF1A alleles with digital PCR indicated the presence of both methylated and unmethylated *RASSF1A* alleles in two of these commercial control samples (**Figure 4E, Figure S1**). Consequently, these samples cannot be used as absolute references in quantitative experiments. For MS-MCA they may fit, as they provide control melting peaks that are useful for a qualitative characterization. Still, the use of these controls to correct for non-specific and incomplete DNA digestion should be given careful consideration, as it may introduce a systematic mathematical error in the obtained quantifications.

On the other hand, the limitations of our digital approach should also be taken into account. Although we confirmed theoretically that our mathematical rationale is valid and that the methylation fraction can be correctly determined, our in-silico simulations also indicated that the uncertainty of the results depends on the amount of input DNA and the methylation fraction itself (see **Supplementary data 1** and **Figure 3**). Whereas the input may be adjusted experimentally and its varying influence on result precision is a known characteristic of digital PCR, the methylation fraction is sample-dependent and cannot be changed (14). Generally, lower input concentrations and higher methylation fractions are accompanied by a lower absolute uncertainty. However, as the digital error of the experiments can be correctly estimated, this uncertainty is readily translated into broader confidence intervals (see **Figure 3C and D**). These findings were confirmed and clearly illustrated by our calibration curve experiments (see **Figure 6B**).

Moreover, the approach described in this manuscript is not specifically designed for rare allele detection. Instead, we validated that our setup is providing accurate quantifications of methylation fractions across the whole range from 0% to 100% (**Figure 6B**). This is comparable to the application of digital PCR in DNA-based T-cell quantifications and covers the biological relevance of DNA methylation in most sample mixtures of hyper- or hypomethylated alleles (5).

Finally, our experimentally prepared mixtures provide a proof of concept of an extended multiplex setup of our approach to determine the allele-specificity of the DNA methylation (see **Figure 7D**). While a heterozygous SNP nearby the DNA methylation target region serves as allelic reference, our target assay and MSRE incubation indicate which alleles are methylated. When intact alleles are partitioned into the droplets, the (lack of) double-positivity (i.e. linkage) for a SNP variant and DNA methylation target is a quantitative measure of the allele-specificity. We showed that this multiplex setup can be effectively applied to calculate the initial linkage (i.e. the baseline conditions, how many of all alleles are intact and not fragmented), and – in combination with a MSRE – to assess the allele-specificity of DNA methylation.

Whereas other approaches determine allele-specificity in haplotype-preserving long PCR experiments, our setup only relies on the intactness of alleles during sample partitioning (24). The number of intact alleles is measurable in the *ASM/Baseline* experiment without a MSRE, while the *ASM/MSRE+* experiment specifies the analysis to the methylated alleles. Following the same experimental rationale, specific interactions between genetic (SNP’s, mutations) and epigenetic alterations (absence or presence of DNA methylation) may now be studied in a quantitative manner.

To conclude, our results demonstrate the possibilities of accurately quantifying DNA methylation with digital PCR, independent of sodium bisulphite conversion. A superior measurement precision could be obtained compared to traditional techniques, and without the disadvantages of the conversion, more effective experiments may be carried out.

Using digital PCR, a targeted density of DNA methylation can now be quantified comparably to genetic alterations as mutations or copy number alterations. We therefore propose that (sub) clone- or cell-type-specific DNA methylation markers, such as, but not limited to *RASSF1A*, may be investigated accordingly in both benign and malignant samples, possibly providing new insights in both human health and disease. Moreover, as the context-density and allele-specificity of this DNA methylation markers can also be determined, important biological mechanisms can now be quantitatively assessed in a digital experiment.

## ACKNOWLEDGEMENTS

We thank our colleagues from the Department of Immunohematology and Blood Transfusion, Leiden University Medical Center, Leiden, the Netherlands) for providing PBMC and placental reference samples, N.A. Gruis (Department of Dermatology, Leiden University Medical Center, Leiden, the Netherlands) for reviewing the manuscript and W.H. Zoutman (Department of Dermatology, Leiden University Medical Center, Leiden, the Netherlands) for helpful discussions and his expertise regarding sodium bisulphite experiments.

## AVAILABILITY

- R Core Team (2018). R: A language and environment for statistical computing. R Foundation for Statistical Computing, Vienna, Austria. https://www.R-project.org/.
- JJ Allaire et al. (2018). rmarkdown: Dynamic Documents for R. https://rmarkdown.rstudio.com.
- RJ Nell et al. (2019). digitalPCRsimulations: In silico digital PCR for R. https://git.lumc.nl/rjnell/digitalpcrsimulations.

## SUPPLEMENTARY

**Supplementary table 1.**
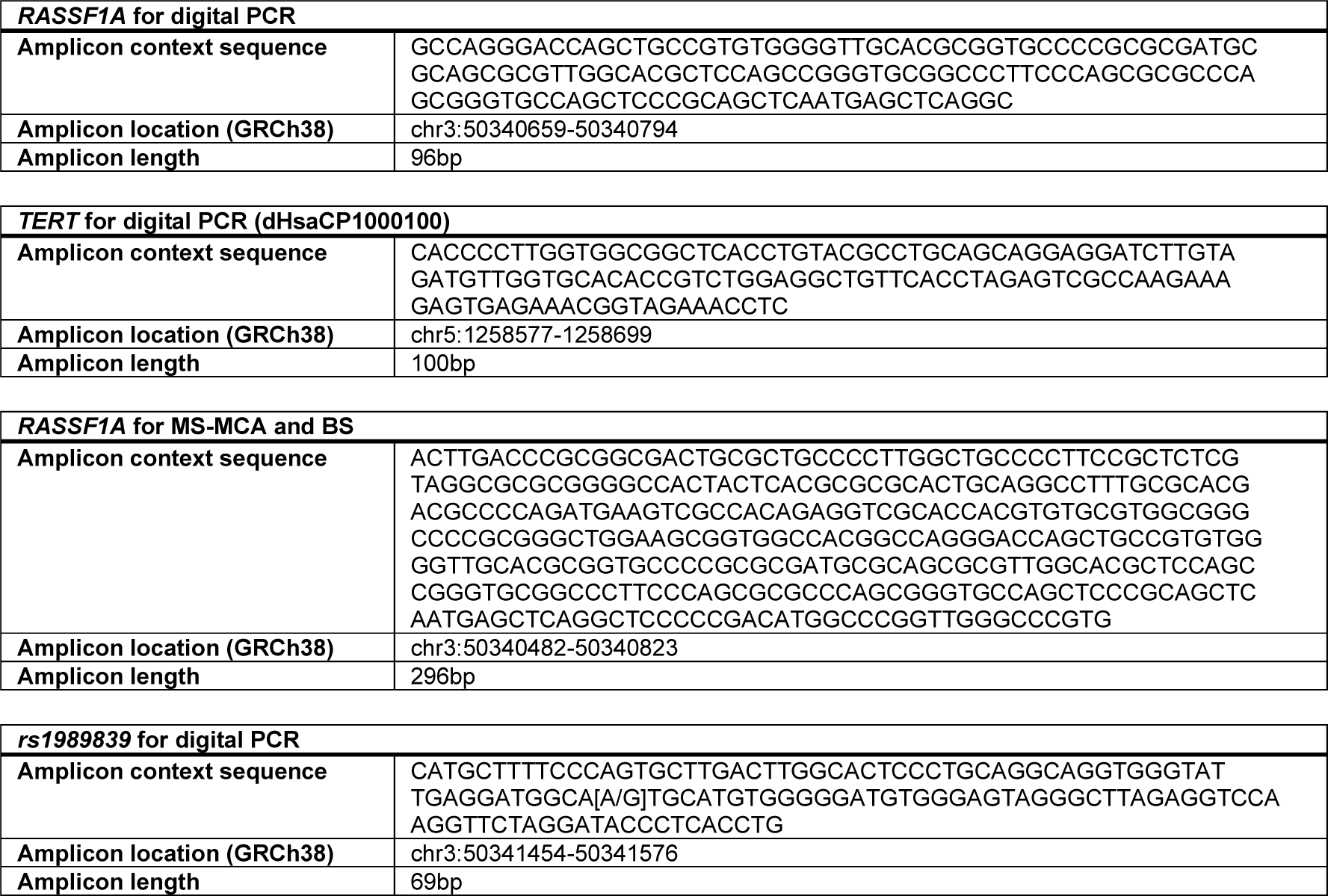
MIQE sequences for digital PCR, MS-MCA and BS assays.

**Supplementary figure 1.**
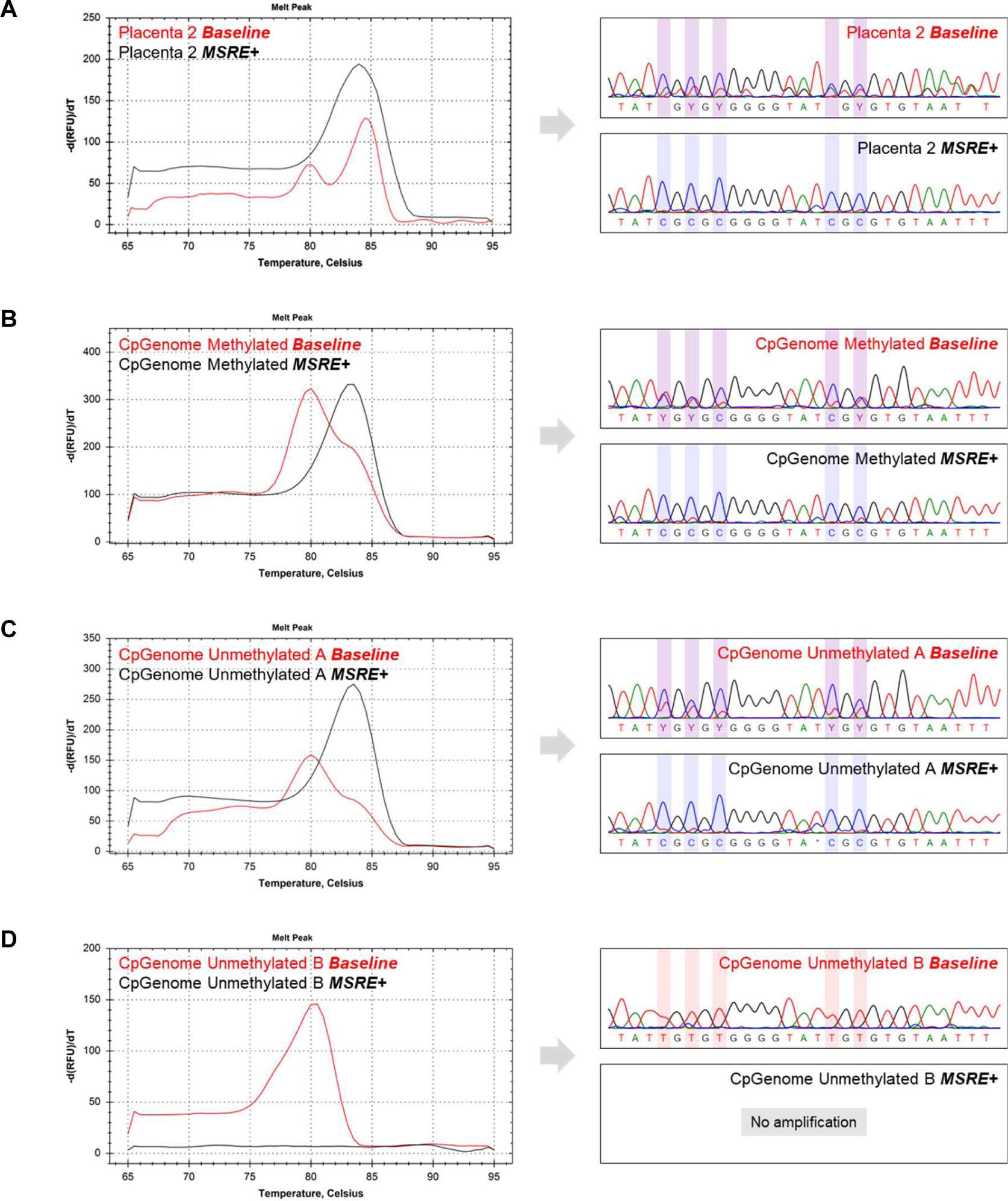
MS-MCA and BS results for sodium bisulphite converted DNA of additional reference samples. **(A)** Partially methylated Placenta 2. **(B)** Partially methylated CpGenome Methylated. **(C)** Partially methylated CpGenome Unmethylated A. **(D)** Unmethylated CpGenome Unmethylated B.

**Supplementary data 1** Overview of in-silico simulation of experimental setup to validate mathematical approach.

**Supplementary data 2** Overview of mathematical derivation to calculate linkage in multiplex digital PCR and in-silico simulation of experimental setup to validate this approach.

