## Supplementary data 1 for "Quantification of DNA methylation using methylation-sensitive restriction enzymes and multiplex digital PCR"

*Nell et al. - Quantification of DNA methylation using methylation-sensitive restriction enzymes and multiplex digital PCR*

#### Background

In this document, the in silico simulation of our digital PCR approach to quantify methylation is described. Moreover, the results of two simulation runs are presented. The same scripts are used to generate *Figure 3* in the original manuscript.

### In silico simulation

#### Dependencies

To simulate the digital PCR experiments, we use our R library `digitalPCRsimulations`, which can be found here: https://git.lumc.nl/rjnell/digitalPCRsimulations.

```
# Load library
library(digitalPCRsimulations)

# Set random seed
set.seed(12345)
```

#### Function `methylation.simulation`

*Baseline* and *MSRE+* are simulated and the obtained ratios are combined to calculate the methylation fraction. The mathematical rationale is outlined in the *Methods* section of the original manuscript.

```
methylation.simulation = function(input_ng, methylation_fraction, n_droplets, n_simulations, alpha=0.95) {

  # Baseline experiment

    # Create universes
    baseline.universe_target = universe(input_ng)
    baseline.universe_reference = universe(input_ng)

    # Simulate ratio experiments
    baseline.ratios = simulate_ratios(baseline.universe_target,
                                      baseline.universe_reference,
                                      n_droplets,
                                      n_simulations,
                                      alpha)

  # MSRE+ experiment

    # Create universes
    msre.universe_target = universe(input_ng * methylation_fraction)
    msre.universe_reference = universe(input_ng)

    # Simulate ratio experiments
    msre.ratios = simulate_ratios(msre.universe_target,
                                  msre.universe_reference,
                                  n_droplets,
                                  n_simulations,
                                  alpha)

  # Simulate the methylation.ratio experiments
  methylation.ratios = NULL
  for (sim_n in 1:n_simulations) {
    methylation.ratios = rbind(methylation.ratios, calc_ratio(msre.ratios[sim_n,], baseline.ratios[sim_n,]))
  }

  # Return simulation result
  return(methylation.ratios)

}
```

### Simulations

#### Simulation 1

##### Building the simulation

In this first simulation the following parameters are evaluated:

- 20 ng genomic input DNA;
- a methylation fraction of 50%;
- 20,000 droplets;
- 10,000 in silico experiments;
- a 95% level of significance.

```
simulation_1 = methylation.simulation(input_ng = 20,
                                      methylation_fraction = .5,
                                      n_droplets = 20000,
                                      n_simulations = 10000,
                                      alpha = .95)
```

##### Summary of the results

The results of the 50 first experiments are sorted and plotted:

```r
par(mar=c(5.1, 5.1, 4.1, 2.1))
plot_simulations(results = simulation_1[1:50,],
                 true_value = .5,
                 error = T,
                 main = "Simulation 1")
```

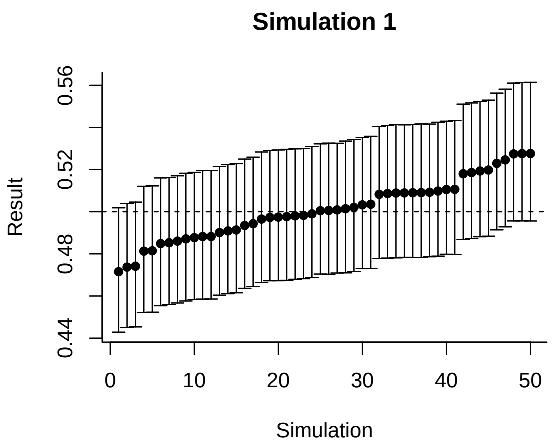

This plot demonstrates that the calculated methylation fractions have a point estimate mean around our true methylation fraction of 50%. Moreover, the calculated 95% confidence intervals generally contain this true methylation fraction.

Based on all simulations, the following statistics are calculated:

```r
stats(simulation_1, true_value = 0.5)
```

```
## $not_in_interval
## [1] 499
##
## $in_interval
## [1] 9501
##
## $coverage
## [1] 0.9501
##
## $point_estimate_mean
## [1] 0.5003596
##
## $point_estimate_sd
## [1] 0.01588745
```

The 95% CI-coverage (i.e. the percentage calculated 95% confidence intervals that contained the true methylation fraction) is ~**95%.** This is close to our intended 95%, demonstrating the mathematical correctness of confidence interval calculation in these simulated conditions.

#### Simulation 2

##### Building the simulation

In this second simulation a range of 20 methylation fractions from 5% to 100% is evaluated:

- 20 ng genomic input DNA;
- 20 methylation fractions (5%, 10%, . . . , 95%, 100%);
- 20,000 droplets;
- 1,000 in silico experiments;
- a 95% level of significance.

```
simulation_2 = list()

# Define range of methylation fractions to simulate
methylation_fractions = seq(from = 0.05,
                            to = 1,
                            by = 0.05)

# Iterate through methylation fractions
for (methylation_fraction in methylation_fractions) {

  # Perform simulation and save results
  simulation_2[[paste(methylation_fraction)]] =
    methylation.simulation(input_ng = 20,
                           methylation_fraction = methylation_fraction,
                           n_droplets = 20000,
                           n_simulations = 1000,
                           alpha = .95)
}
```

##### Summary of the results

The results of the first experiment of each methylation fraction are plotted:

```
x = methylation_fractions
y = NULL
col = NULL
for (methylation_fraction in methylation_fractions) {
  res = simulation_2[[paste(methylation_fraction)]][1,]
  y = rbind(y, res)
  if (res[3] < methylation_fraction | res[2] > methylation_fraction) {
    col = c(col, "red")
  }
  else {
    col = c(col, "black")
  }
}

par(mar=c(5.1, 5.1, 4.1, 2.1))
plot(x = methylation_fractions,
     y = y[,1],
     xlab = "Simulated methylation fraction",
     ylab = "Calculated methylation fraction",
     xlim = c(0, 1.05), ylim = c(0, max(y)),
     bty = "l", axes = F, xaxs = "i", yaxs = "i", type = "n",
     main = "Simulation 2 - First exploration")
grid(col="#eeeeee", lty=1)
segments(0,0,1.05,0,xpd=T)
segments(0,0,0,max(y),xpd=T)
axis(side = 2, las=1)
axis(side = 1, las=1)
segments(0.05, 0.05, 1, 1, lty=2)
arrows(x, y[,2], x, y[,3], length=0.05, angle=90, code=3, col=col, xpd=T)
points(x = x, y = y[,1], pch = 16, xpd = T)
```

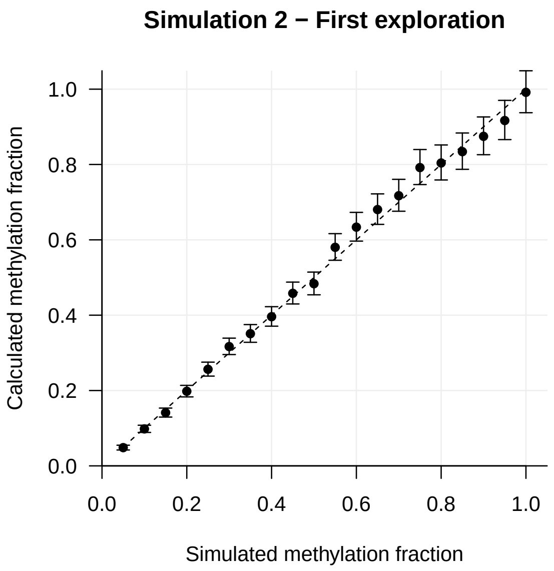

This plot demonstrates that all calculated methylation fractions (y-axis) have a point estimate around the true methylation fraction (x-axis). Moreover, the calculated 95% confidence intervals generally contain this true methylation fraction. The absolute width of the confidence intervals is typically larger at higher methylation fractions.

Based on all simulations per methylation fraction, the statistics are calculated and stored:

```r
# Initialize result vectors
means = NULL
sds = NULL
coverages = NULL
widths = NULL

# Iterate through methylation fractions
for (methylation_fraction in methylation_fractions) {

  # Calculate stats
  this_stats = stats(simulation_2[[paste(methylation_fraction)]], true_value = methylation_fraction)

  # Save results
  means = c(means, this_stats$point_estimate_mean)
  sds = c(sds, this_stats$point_estimate_sd)
  coverages = c(coverages, this_stats$coverage)
  widths = c(widths, mean(simulation_2[[paste(methylation_fraction)]][,3] -
                            simulation_2[[paste(methylation_fraction)]][,2]))
}
```

##### Mean per methylation fraction

The mean of all simulations per methylation fraction is plotted:

```
par(mar=c(5.1, 5.1, 4.1, 2.1))
plot(x = methylation_fractions,
     y = means,
     xlab = "Simulated methylation fraction",
     ylab = "Calculated methylation fraction",
     xlim = c(0,1), ylim = c(0,1),
     bty = "l", axes = F, xaxs = "i", yaxs = "i", type = "n",
     main = "Simulation 2 - Means")
grid(col="#eeeeee", lty=1)
axis(side = 2, las=1)
axis(side = 1, las=1)
segments(0.05, 0.05, 1, 1, lty=2)
points(x = x, y = means, pch = 16, xpd = T)
```

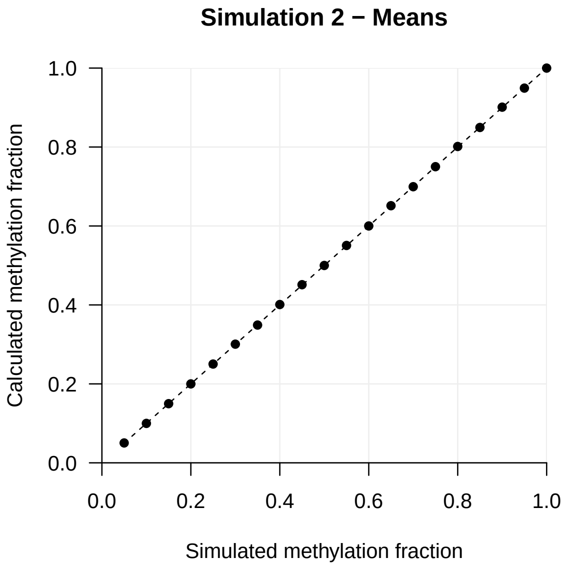

This shows an excellent correlation between calculated methylation fraction (y-axis) and true methylation fraction (x-axis), indicating the underlying mathematical approach is valid.

##### Standard deviation per methylation fraction

The standard deviation per methylation fraction is plotted:

```
par(mar=c(5.1, 5.1, 4.1, 2.1))
plot(x = methylation_fractions,
     y = sds,
     xlab = "Simulated methylation fraction",
     ylab = "",
     xlim = c(0,1), ylim = c(0,max(sds)),
     bty = "l", axes = F, xaxs = "i", yaxs = "i", type = "n",
     main = "Simulation 2 - Standard deviations")
grid(col="#eeeeee", lty=1)
title(ylab="Standard deviation", line=4)
segments(0,0,0,max(sds),xpd=T)
axis(side = 2, las=1)
axis(side = 1, las=1)
points(x = x, y = sds, pch = 16, xpd = T)
```

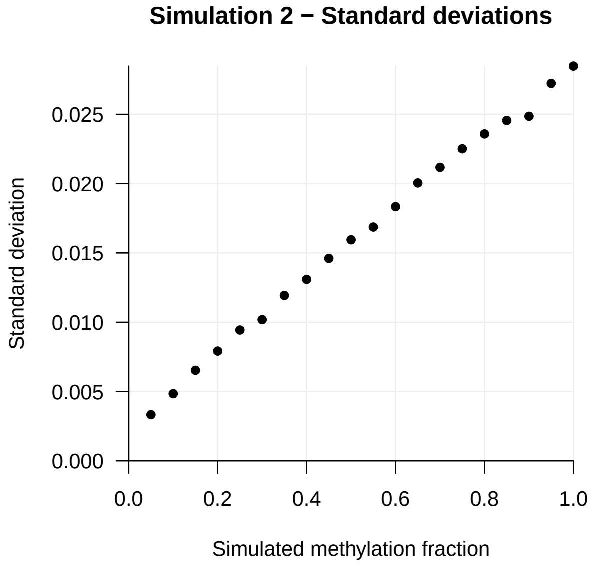

Larger standard deviations are found when the true methylation fraction is higher. This was already suggested by the earlier observation that higher methylation fractions are accompanied by larger confidence intervals. In conclusion, the absolute uncertainty is larger at a higher methylation fraction.

##### 95% CI-coverage per methylation fraction

The 95% CI-coverage (i.e. the percentage calculated 95% confidence intervals that contained the true methylation fraction) per methylation fraction is plotted:

```
par(mar=c(5.1, 5.1, 4.1, 2.1))
plot(x = methylation_fractions,
     y = coverages,
     xlab = "Simulated methylation fraction",
     ylab = "95% CI-coverage",
     xlim = c(0,1), ylim = c(0,1),
     bty = "l", axes = F, xaxs = "i", yaxs = "i", type = "n",
     main = "Simulation 2 - Coverages")
grid(col="#eeeeee", lty=1)
axis(side = 2, las=1)
axis(side = 1, las=1)
segments(0, 0.95, 1, 0.95, lty=2)
points(x = x, y = coverages, pch = 16, xpd = T)
```

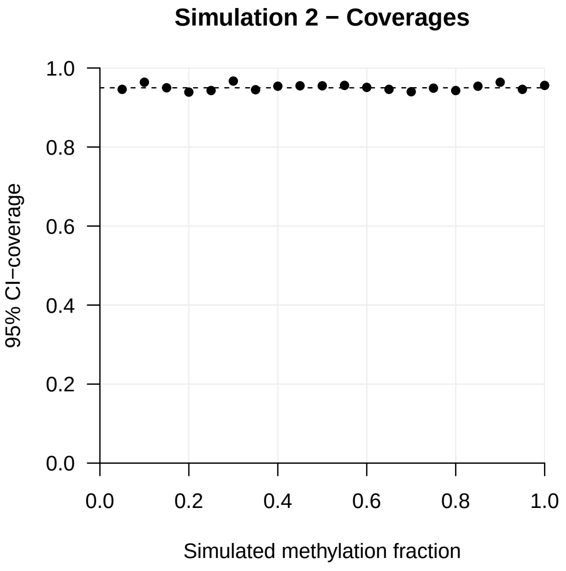

In all simulated methylation fractions this coverage is close to our intended 95%, demonstrating the mathematical correctness of the confidence interval calculations.

##### Absolute 95% CI-width per methylation fraction

The average absolute width of the 95% confidence intervals per methylation fraction is plotted:

```
par(mar=c(5.1, 5.1, 4.1, 2.1))
plot(x = methylation_fractions,
     y = widths,
     xlab = "Simulated methylation fraction",
     ylab = "",
     xlim = c(0,1), ylim = c(0,max(widths)),
     bty = "l", axes = F, xaxs = "i", yaxs = "i", type = "n",
     main = "Simulation 2 - Absolute 95% CI-widths")
grid(col="#eeeeee", lty=1)
title(ylab="Absolute 95% CI-width", line=4)
segments(0,0,0,max(widths),xpd=T)
axis(side = 2, las=1)
axis(side = 1, las=1)
points(x = x, y = widths, pch = 16, xpd = T)
```

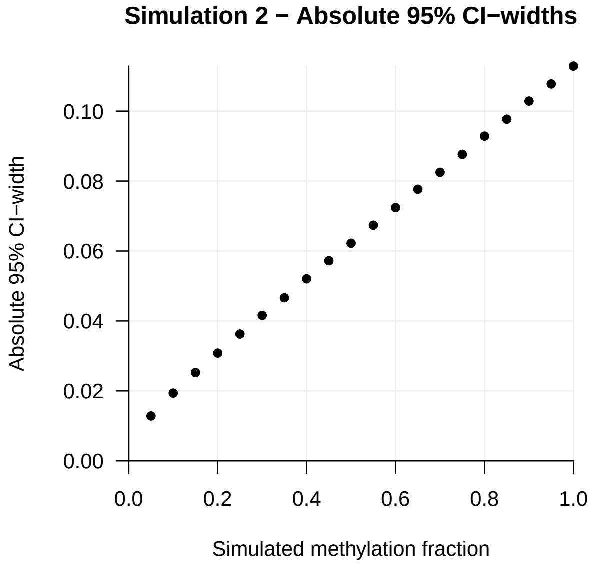

The absolute width of the 95% confidence intervals increases with higher methylation fractions. This relates to the earlier observations that at higher methylation fractions, a larger uncertainty is encountered. However, as the digital error is correctly estimated across all fractions (indicated by the 95% CI-coverage), broader confidence intervals are observed at the less certain higher methylation fractions.
