## Supplementary data 2 for "Quantification of DNA methylation using methylation-sensitive restriction enzymes and multiplex digital PCR"

*Nell et al. - Quantification of DNA methylation using methylation-sensitive restriction enzymes and multiplex digital PCR*

#### Background

In this document, the mathematical formulae for the determination of allele-specific methylation are described. Moreover, an in-silico simulation of the experimental setup is built to validate this approach.

#### Mathematical derivation

The experimental setup of our ASM involves the multiplex determination of SNP1, SNP2 and target concentrations simultaneously. This multiplex can be seen as the sum of 5 'virtual' experiments, in which specific alleles (linked double-positive [i.e. *SNP1-T* or *SNP2-T*] or unlinked single-positive [i.e. *SNP1**, *SNP2** or *T**]) are distributed among the droplets (see *Figure 1*). When in the summed experiment all alleles are mixed and measured together in the multiplex setup, randomly droplets will become double or triple positive, as some alleles will copartition. Generally, this will result in single-positive clusters to become smaller.

The proportion empty droplets $p_1$ in the true multiplex can be defined as the following product of virtual proportions:

$$p_1 = (1 - p_{SNP1^*}) \cdot (1 - p_{T^*}) \cdot (1 - p_{SNP1\text{-}T}) \cdot (1 - p_{SNP2^*}) \cdot (1 - p_{SNP2\text{-}T}) \quad (1)$$

The final proportion of droplets within a single-positive cluster depends on not being copartitioned with another allele:

$$p_2 = p_{SNP1^*} \cdot (1 - p_{T^*}) \cdot (1 - p_{SNP1\text{-}T}) \cdot (1 - p_{SNP2^*}) \cdot (1 - p_{SNP2\text{-}T}) \quad (2)$$
$$p_3 = (1 - p_{SNP1^*}) \cdot p_{T^*} \cdot (1 - p_{SNP1\text{-}T}) \cdot (1 - p_{SNP2^*}) \cdot (1 - p_{SNP2\text{-}T}) \quad (3)$$
$$p_5 = (1 - p_{SNP1^*}) \cdot (1 - p_{T^*}) \cdot (1 - p_{SNP1\text{-}T}) \cdot p_{SNP2^*} \cdot (1 - p_{SNP2\text{-}T}) \quad (4)$$

As many factors of these products are shared, $p_{SNP1}$, the virtual proportion of droplets filled with SNP1 alleles, can be resolved easily:

$$\frac{p_2}{p_2 + p_1} = \frac{p_{SNP1^*} \cdot (1 - p_{T^*}) \cdot (1 - p_{SNP1\text{-}T}) \cdot (1 - p_{SNP2^*}) \cdot (1 - p_{SNP2\text{-}T})}{(p_{SNP1^*} + (1 - p_{SNP1^*})) \cdot (1 - p_{T^*}) \cdot (1 - p_{SNP1\text{-}T}) \cdot (1 - p_{SNP2^*}) \cdot (1 - p_{SNP2\text{-}T})} \quad (5)$$
$$= \frac{p_{SNP1^*}}{p_{SNP1^*} + (1 - p_{SNP1^*})} = \frac{p_{SNP1^*}}{1} = p_{SNP1^*} \quad (6)$$

We consider

$$\hat{p}_{SNP1^*} = \frac{p_2}{p_2 + p_1} \quad (7)$$

as estimator of $p_{SNP1^*}$.

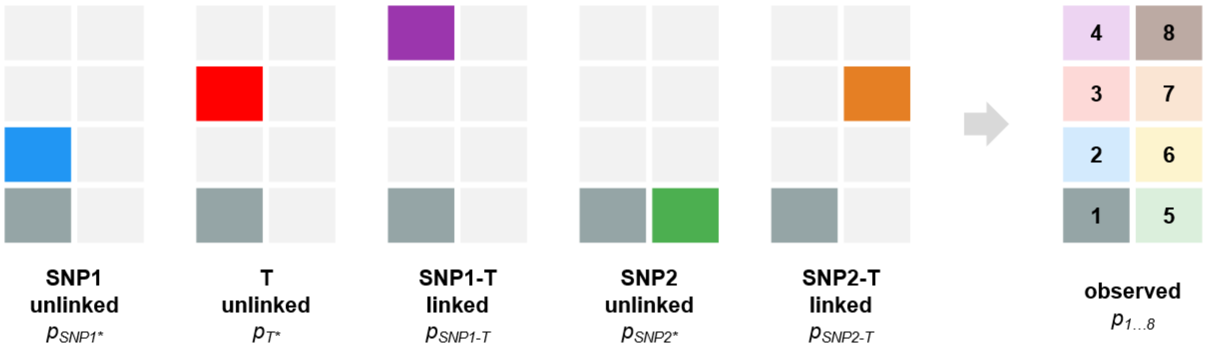

Figure 1: The observed $p_{1...8}$ from the multiplex setup are derived from the combination of 5 'virtual' experiments.

Following the work of Dube et al. (2008), the true concentration of $\lambda$ of SNP1* molecules can be estimated by:

$$\hat{\lambda}_{SNP1^*} = -\ln(1 - \hat{p}_{SNP1^*}) \tag{8}$$

When assuming

$$\hat{\lambda}_{SNP1} = \hat{\lambda}_{SNP1^*} + \hat{\lambda}_{SNP1\text{-}T} \tag{9}$$

the linkage $\Lambda_{SNP1}$, the fraction linked alleles of SNP1, can be estimated as follows:

$$\hat{\Lambda}_{SNP1} = 1 - \frac{\hat{\lambda}_{SNP1^*}}{\hat{\lambda}_{SNP1}} \tag{10}$$

### In silico simulation

#### Dependencies

To simulate the digital PCR experiments, we use our R library `digitalPCRsimulations`, which can be found here: https://git.lumc.nl/rjnell/digitalPCRsimulations.

```
# Load library
library(digitalPCRsimulations)

# Set random seed
set.seed(12345)
```

To evaluate the linkage-calculations of our simulations, we create two additional functions:

- `merge_universes(...)` to combine several universes into one complete universe.
- `calc_conc_unlinked(...)` to calculate the concentration unlinked from the given universes.

```
merge_universes = function (u1, u2, u3=NULL) {
  u2[which(u1 == 1)] = 1
  if (!is.null(u3)) {
    u2[which(u3 == 1)] = 1
  }
  return(u2)
}
```

```
calc_conc_unlinked = function(u1, u2, u3, sample, alpha = 0.95) {

  u1 = u1[sample]
  u2 = u2[sample]
  u3 = u3[sample]

  # Define the sample
  count = length(which(u1 == 1 & u2 == 0 & u3 == 0))
  droplets = length(which(u1 == 0 & u2 == 0 & u3 == 0)) + count

  p = count / droplets

  # Translate this percentage to concentration...
  conc = -log(1-p)

  # Return the calculated concentration
  return(conc/0.00085)
}
```

#### Function `linkage.simulation`

An *in-silico* linkage experiment is carried out, a setup outlined in the *Methods* section of the original manuscript.

```
linkage.simulation = function(input_ng, linkage_SNP1, linkage_SNP2, n_droplets, n_simulations, alpha=0.95) {

  # Create separate universe for SNP1 unlinked alleles
  universe_SNP1 = universe(input_ng/2 * (1-linkage_SNP1))

  # Create separate universe for T-SNP1 linked alleles
  universe_T_SNP1 = universe(input_ng/2 * linkage_SNP1)

  # Create separate universe for SNP2 unlinked alleles
  universe_SNP2 = universe(input_ng/2 * (1-linkage_SNP2))

  # Create separate universe for T-SNP2 linked alleles
  universe_T_SNP2 = universe(input_ng/2 * linkage_SNP2)

  # Create separate universe for T-SNP2 linked alleles
  universe_T = universe(input_ng/2 * (1-linkage_SNP1) + input_ng/2 * (1-linkage_SNP2))

  # As all alleles are mixed in the final multiplex experiment,
  # the separate universes are merged:
  multiplex.universe_SNP1 = merge_universes(universe_SNP1, universe_T_SNP1)
  multiplex.universe_SNP2 = merge_universes(universe_SNP2, universe_T_SNP2)
  multiplex.universe_T = merge_universes(universe_T_SNP1, universe_T_SNP2, universe_T)
  multiplex.universe_SNP12 = merge_universes(multiplex.universe_SNP1, multiplex.universe_SNP2)

  # Initialize results vectors
  results_linkage_SNP1 = NULL
  results_linkage_SNP2 = NULL

  for (simulation in 1:n_simulations) {

    # Simulate a sample to obtain from the universes
    sample = simulate_sample(n_droplets)

    # Calculate concentration total SNP1
    concentration_SNP1 = sample_from_universe(multiplex.universe_SNP1,
                                              sample,
                                              alpha)

    # Calculate concentration unlinked SNP1
    concentration_SNP1_unlinked = calc_conc_unlinked(multiplex.universe_SNP1,
                                                     multiplex.universe_SNP2,
                                                     multiplex.universe_T,
                                                     sample,
                                                     alpha)

    # Calculate linkage SNP1
    linkage_SNP1 = 1 - concentration_SNP1_unlinked / concentration_SNP1[1]

    # Save result
    results_linkage_SNP1 = rbind(results_linkage_SNP1, linkage_SNP1)

    # Calculate concentration total SNP2
    concentration_SNP2 = sample_from_universe(multiplex.universe_SNP2,
                                              sample,
                                              alpha)

    # Calculate concentration unlinked SNP2
    concentration_SNP2_unlinked = calc_conc_unlinked(multiplex.universe_SNP2,
                                                     multiplex.universe_SNP1,
                                                     multiplex.universe_T,
                                                     sample,
                                                     alpha)
```

```
    # Calculate linkage SNP2
    linkage_SNP2 = 1 - concentration_SNP2_unlinked / concentration_SNP2[1]

    # Save result
    results_linkage_SNP2 = rbind(results_linkage_SNP2, linkage_SNP2)

  }

  return(cbind(results_linkage_SNP1, results_linkage_SNP2))
}
```

### Simulations

#### Simulation 1

##### Building the simulation

In this first simulation the following parameters are evaluated:

- 20 ng genomic input DNA;
- heterozygous, 50% SNP1 and 50% SNP2;
- 50% linkage for SNP1;
- 0% linkage for SNP2;
- 20,000 droplets;
- 1,000 in silico experiments;
- a 95% level of significance.

```
simulation_1 = linkage.simulation(input_ng = 20,
                                  linkage_SNP1 = 0.5,
                                  linkage_SNP2 = 0,
                                  n_droplets = 20000,
                                  n_simulations = 1000,
                                  alpha = .95)
```

##### Summary of the results

The mean linkage for SNP1 and SNP2 as determined by the simulations are calculated:

```
# Mean linkage SNP1
mean(simulation_1[,1])
```

```
## [1] 0.4979186
```

The mean determined linkage for SNP1 is ~**49.8%**, which is close to the true linkage of 50%, demonstrating the correctness of the mathematical approach under these simulated conditions.

```
# Mean linkage SNP2
mean(simulation_1[,2])
```

```
## [1] -0.0001433718
```

The mean determined linkage for SNP2 is ~**0%**, which is close to the true linkage of 0%, demonstrating the correctness of the mathematical approach under these simulated conditions.

#### Simulation 2

##### Building the simulation

In this second simulation the following parameters are evaluated:

- 20 ng genomic input DNA;
- heterozygous, 50% SNP1 and 50% SNP2;
- 20 linkage values for for SNP1 (5%, 10%, . . . , 95%, 100%);
- 20 linkage values for for SNP2, inverse to SNP1 (100%, 95%, . . . , 10%, 5%);
- 20,000 droplets;
- 1,000 in silico experiments;
- a 95% level of significance.

```
simulation_2 = list()

# Define range of methylation fractions to simulate
linkages = seq(from = 0.05,
               to = 1,
               by = 0.05)

# Iterate through methylation fractions
for (linkage in linkages) {

  # Perform simulation and save results
  simulation_2[[paste(linkage)]] =
    linkage.simulation(input_ng = 20,
                       linkage_SNP1 = linkage,
                       linkage_SNP2 = 1-linkage,
                       n_droplets = 20000,
                       n_simulations = 1000,
                       alpha = .95)
}

# Initialize result vectors
means = NULL
sds = NULL

# Iterate through methylation fractions
for (linkage in linkages) {

  # Save results
  means = c(means, colMeans(simulation_2[[paste(linkage)]])[1])
  sds = c(sds, matrixStats::colSds(simulation_2[[paste(linkage)]])[1])
}
```

##### Mean per simulated linkage value

The mean of all simulations per simulated linkage value is plotted:

```
par(mar=c(5.1, 5.1, 4.1, 2.1))
plot(x = linkages,
     y = means,
     xlab = "Simulated linkage",
     ylab = "Calculated linkage",
     xlim = c(0,1), ylim = c(0,1),
     bty = "l", axes = F, xaxs = "i", yaxs = "i", type = "n",
     main = "Simulation 2 - Means")
grid(col="#eeeeee", lty=1)
axis(side = 2, las=1)
axis(side = 1, las=1)
segments(0.05, 0.05, 1, 1, lty=2)
points(x = linkages, y = means, pch = 16, xpd = T)
```

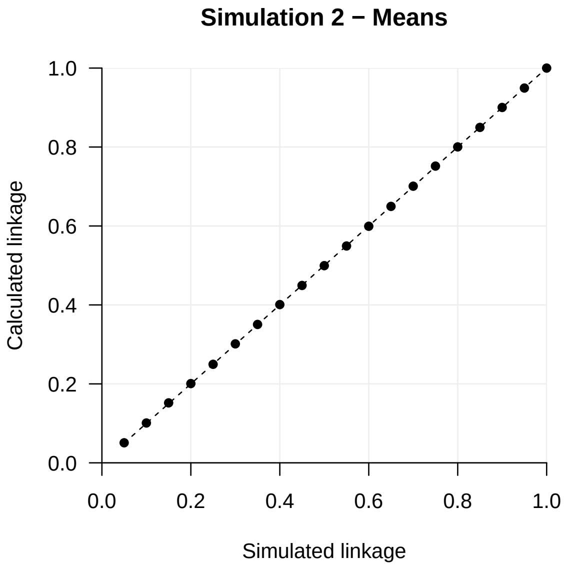

This shows an excellent correlation between calculated linkage (y-axis) and true linkage (x-axis), indicating the underlying mathematical approach is valid.

##### Standard deviation per simulated linkage value

The standard deviation per simulated linkage value is plotted:

```
par(mar=c(5.1, 5.1, 4.1, 2.1))
plot(x = linkages,
     y = sds,
     xlab = "Simulated linkage",
     ylab = "",
     xlim = c(0,1), ylim = c(0,max(sds)),
     bty = "l", axes = F, xaxs = "i", yaxs = "i", type = "n",
     main = "Simulation 2 - Standard deviations")
grid(col="#eeeeee", lty=1)
title(ylab="Standard deviation", line=4)
segments(0,0,0,max(sds),xpd=T)
axis(side = 2, las=1)
axis(side = 1, las=1)
points(x = linkages, y = sds, pch = 16, xpd = T)
```

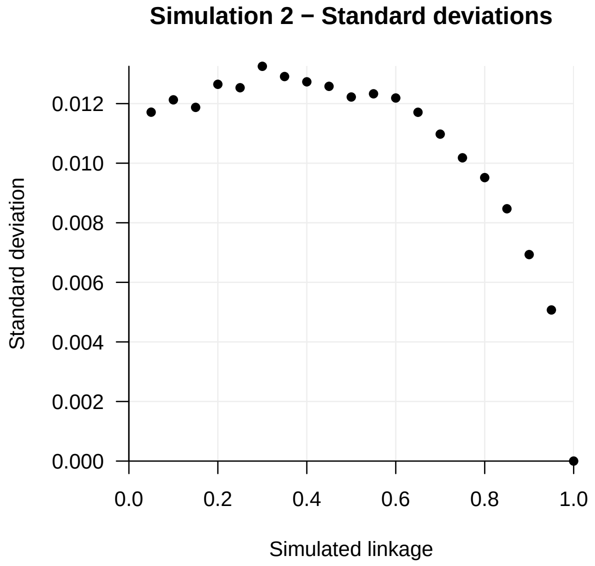

Larger standard deviations are found when the true linkage is lower, but all in all the uncertainty is rather limited.
